# High throughput single cell RNA-seq of developing mouse kidney and human kidney organoids reveals a roadmap for recreating the kidney

**DOI:** 10.1101/235499

**Authors:** Alexander N. Combes, Belinda Phipson, Luke Zappia, Kynan T. Lawlor, Pei Xuan Er, Alicia Oshlack, Melissa H. Little

**Author notes:** Equal contribution (Alexander N. Combes, Belinda Phipson). Corresponding authors M.H.L.; A.N.C.

## Abstract

Recent advances in our capacity to differentiate human pluripotent stem cells to human kidney tissue are moving the field closer to novel approaches for renal replacement. Such protocols have relied upon our current understanding of the molecular basis of mammalian kidney morphogenesis. To date this has depended upon population based-profiling of non-homogenous cellular compartments. In order to improve our resolution of individual cell transcriptional profiles during kidney morphogenesis, we have performed 10x Chromium single cell RNA-seq on over 6000 cells from the E18.5 developing mouse kidney, as well as more than 7000 cells from human iPSC-derived kidney organoids. We identified 16 clusters of cells representing all major cell lineages in the E18.5 mouse kidney. The differentially expressed genes from individual murine clusters were then used to guide the classification of 16 cell clusters within human kidney organoids, revealing the presence of distinguishable stromal, endothelial, nephron, podocyte and nephron progenitor populations. Despite the congruence between developing mouse and human organoid, our analysis suggested limited nephron maturation and the presence of ‘off target’ populations in human kidney organoids, including unidentified stromal populations and evidence of neural clusters. This may reflect unique human kidney populations, mixed cultures or aberrant differentiation *in vitro*. Analysis of clusters within the mouse data revealed novel insights into progenitor maintenance and cellular maturation in the major renal lineages and will serve as a roadmap to refine directed differentiation approaches in human iPSC-derived kidney organoids.

## Introduction

Knowledge of developmental programs can be used to direct the differentiation of human induced pluripotent stem cells towards a desired cell fate. Such approaches have successfully generated models of human intestinal epithelium, brain, and ear, in each instance forming multicellular self-organising structures termed organoids by mimicking conditions that regulate development of the same tissues during embryogenesis (McCauley and Wells, 2017). Similarly, protocols for the generation of human kidney cells types have been developed by ourselves (Takasato et al., 2014; Takasato et al., 2016b) and others (Freedman et al., 2015; Morizane et al., 2015; Taguchi et al., 2014; Taguchi and Nishinakamura, 2017). Our protocol is based on a stepwise recapitulation of mouse embryogenesis from primitive streak to kidney and results in the formation of kidney organoids containing nephron, stromal and endothelial compartments (Takasato et al., 2014; Takasato et al., 2016b). Such protocols raise the exciting prospect of kidney disease modelling, toxicity and drug screening, and even the production of human kidney cells for novel regenerative medicine applications. However, the utility of kidney tissue generated from stem cells will depend on how faithfully these replicate genuine human kidney morphogenesis and the degree to which the component cell types mature. Our ability to judge the authenticity of engineered tissue depends on our understanding of the markers that define a particular cell type or state of maturation in native tissue. Likewise, capacity to generate a realistic cell type depends on our understanding of the programs that regulate the maintenance of a cellular identity, and the progression from one cell type to another during normal development.

Initial transcriptional comparisons between our human kidney organoids and human development suggested that organoids most closely match trimester 1 human kidney, highlighting the fact that at present these represent models of a developing tissue. Mammalian kidney development has been studied using the mouse for over 70 years. The developing mammalian kidney consists of three main cell lineages, all of which derive from multipotent progenitors. *Foxd1*-expressing stromal progenitors give rise to all cell types in the interstitial compartments (Kobayashi et al., 2014) while *Ret*-expressing ureteric tip (UT) cells give rise to the collecting duct and ureter (Chi et al., 2009). Finally, the filtration units of the kidney, the epithelial nephrons, arise from *Six2*-expressing nephron progenitor cells (Kobayashi et al., 2008). During kidney development, these progenitor populations signal to each other to ensure the ongoing expansion of the organ and accumulation of nephrons, with the resulting organ containing approximately 1,000,000 nephrons in human and 16,000 in mouse (Bertram et al., 2011; Merlet-Benichou et al., 1999). Lineage tracing in mouse has shown that nephrons arise from a mesenchymal nephron progenitor population. These cells undergo a mesenchyme-to-epithelial transition to form renal vesicles, which then undergo elongation, patterning and segmentation to give rise to functionally specialised cell types unique to particular nephron segments. Ultimately, the proximal nephron forms the vascularised glomerulus through which the blood is filtered while the distal end connects to the collecting duct system to allow an exit tract for the urinary filtrate. Specialised tubular cell types form along the intervening nephron segments (proximal tubule, loop of Henle, distal tubule) to facilitate appropriate reabsorption and excretion.

The developing mouse kidney has been intensely analysed at the level of gene expression (www.gudmap.org; (McMahon et al., 2008)), with a tissue atlas initially defined using microarrays of laser-captured cellular compartments followed by section *in situ* hybridisation (Brunskill et al., 2008). While this and subsequent transcriptional studies represented an invaluable resource for the field, most profiled populations were not homogeneous, several cell types are not represented and there are limitations on being able to compare between data sets generated using distinct platforms. More recently, single cell RNA sequencing has resulted in substantial new insight into the cellular composition of complex tissues, including the kidney (Adam et al., 2017; Magella et al., 2017), and has facilitated the discovery of rare cell types and differentiation dynamics within a cell type (Lake et al., 2016).

In this study, we used single cell profiling to define cell types and interrogate signalling pathways within both E18.5 developing mouse kidney and human iPSC-derived kidney organoids. A detailed interrogation of cellular heterogeneity within the developing mouse kidney identified a previously unidentified *Six2*^+^ subpopulation with features of both nephron progenitor and stroma. *In vivo* lineage analysis supported the existence of this population while pseudotime analyses suggested such cells may represent an alternative endpoint for cap mesenchyme and/or an alternative source of nephron cells. Also evident was a transcriptional congruence between distal tubule and ureteric epithelium signatures despite distinct lineage relationships. Correlation between mouse and organoid datasets facilitated an improved understanding of the cell types present within kidney organoids, revealing evidence for mesenchymal, nephron and endothelial compartments equivalent in relative proportion to what is present in the developing mouse. While nephron segmentation was evident and a minor nephron progenitor cluster present in human kidney organoids, profiling revealed the nephron cell types to be immature and proximally-biased with no definitive evidence of collecting duct versus distal tubule / connecting segment. Organoids were also found to contain ‘off-target’ cell populations suggesting either inappropriate differentiation or mixed cultures. New insight into signalling pathways associated with maintenance of cell fate and maturation within the mouse data may offer avenues to improve nephron maturation and refine differentiation protocols for primary mouse progenitors and human pluripotent cells.

## Results

### Single cell profiling of the developing kidney identifies all major lineages and cell types including a novel nephron progenitor state

We first sought to establish a reference data set to explore cell types and developmental programs in the mouse embryonic kidney. We chose 18.5 days of embryonic development (E18.5) because all major embryonic progenitor populations and many maturing cell types co-exist at this time point. Using three independent kidney pairs captured in parallel using the 10x Chromium single cell capture system, our aggregated data set consists of 6752 cells, 5639 of which passed quality control (e.g. proportion of genes with zero read-counts less than 95%), with a median of 2896 unique genes detected per cell. This data has been submitted to NCBI’s Gene Expression Omnibus (Edgar et al, 2002) and is accessible through https://www.ncbi.nlm.nih.gov/geo/query/acc.cgi?acc=GSE108291. We used Seurat (Macosko et al., 2015; Satija et al., 2015), to perform normalisation and unsupervised clustering of cells, settling on an approach that yielded 16 distinct clusters (Fig. 1A, Supplementary Fig. 1AB). The edgeR package was used to test for genes that were differentially expressed between cells in each cluster and all other cells. Genes that were over-expressed or specifically expressed (marker genes) in each cluster, were cross-referenced to validated anchor genes and cell type specific markers to identify cell types (Fig. 1B) (Georgas et al., 2009; Georgas et al., 2008; Thiagarajan et al., 2011b) and entire gene lists were compared to available kidney cell-type specific profiling using ToppGene (toppgene.cchmc.org). This provided a provisional identification for all clusters (Fig. 1A,B). Lists of differentially expressed genes for each cluster are provided in Supplementary Table 1, which features an interactive ‘look up’ sheet, facilitating the interrogation of gene expression in all clusters. One or more clusters representing each of the major renal lineages - stroma, nephron, and ureteric epithelium - were identified in the data (Figure 1A-D). Vascular endothelial and tissue-resident macrophage populations were also identified (Fig. 1A,B). We note that resident macrophages expressed *Bmp2* and *Tgfb1* while the endothelial cells expressed *Tgfb1, Notch1*, and *Notch 2*, which may influence cell-cell signalling within the developing kidney. Clusters corresponding to several distinct nephron progenitor states, all major nephron segments, and a novel nephron progenitor state that co-expresses the definitive nephron progenitor markers *Six2* and *Cited1*, as well as key stromal markers *Pdgfra* and *Col3a1* (Fig. 1A,B) were identified. Four non-nephron progenitor populations with a stromal signature were identified, all expressing *Meis1, Col3a1*, and *Pdgfra*. These populations correspond to cortical (*Foxd1*^+^) and medullary stroma (*Foxd1*^−^), pelvic mesenchyme (*Wnt4*^+^*Wnt11*^+^), and a population that may represent mesangial cells (*Sfrp2*^+^) as previously proposed (Adam et al., 2017). The lists of marker genes and differentially expressed genes for each cluster compare favourably to bulk RNA-seq results from sorted cell populations (O’Brien et al., 2016; Rutledge et al., 2017) while gene by gene analysis of previously identified cell type markers shows anticipated gene expression patterns (Supplementary Table 1). An overlay of the three independent kidneys showed an even distribution of cells from each replicate amongst clusters and a comparison of stage of cell cycle across the t-SNE projection illustrates no association between cell cycle state and any specific cluster after cell cycle normalisation (Supplementary Figure 1AB).

**Figure 1.**
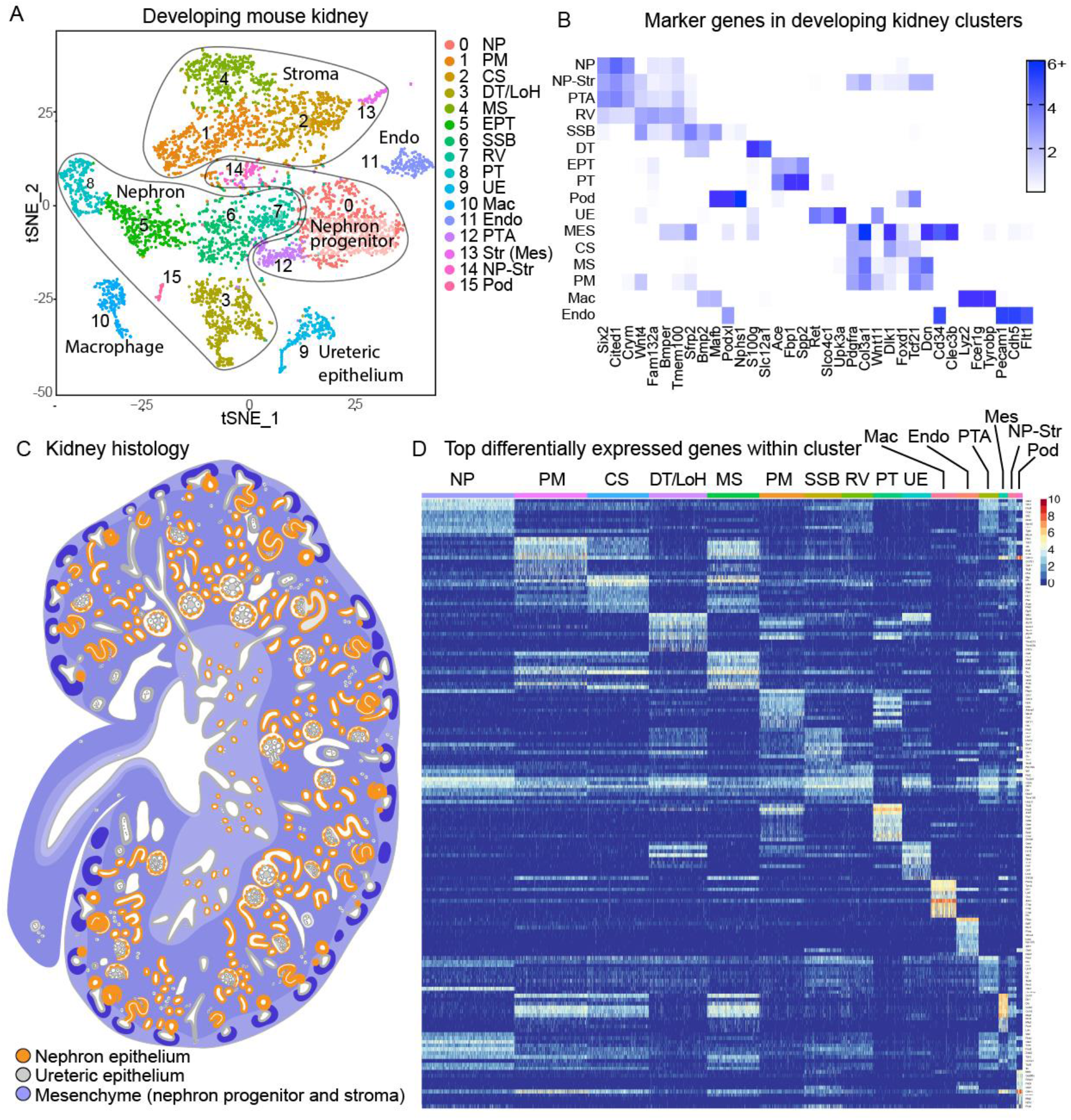
Single cell RNA-seq profiling of the E18.5 mouse kidney reveals 16 distinct cellular clusters. **A**. tSNE plot revealing 16 distinct cell clusters (cluster 0 to cluster 15) identified from largest to smallest population as representing nephron progenitor (NP, cluster 0), pelvic mesenchyme (PM, cluster 1), cortical stroma (CS, cluster 2), distal tubule / loop of Henle (DT/LOH, cluster 3), medullary stroma (MS, cluster 4), early proximal tubule (EPT, cluster 5), S-shaped body (SSB, cluster 6), renal vesicle (RV, cluster 7), proximal tubule (PT, cluster 8), ureteric epithelium (UE, cluster 9), macrophages (Mac, cluster 10), endothelial cells (Endo, cluster 11), pretubular aggregate (PTA, cluster 12), a stromal cluster that may represent mesangial cells (Str (Mes), cluster 13), a novel nephron progenitor – stroma cluster (NP-Str, cluster 14) and podocytes (Pod, cluster 15). **B**. Key differentially expressed genes from each cluster across all 16 cell clusters within the E18.5 developing mouse kidney. Scale indicates log fold change of differential expression relative to all other cells. C. Diagram of the histology of the developing mouse kidney at a similar stage to that analysed illustrating the location of nephron (orange), ureteric epithelial (grey) and mesenchymal (blue) elements within the kidney. Image obtained from the GUDMAP database (gudmap.org/Schematics) D. Heatmap of the top differentially expressed genes across each cluster with cells arranged by cluster from left to right based on descending cluster size.

### Defining the cellular composition of human iPSC-derived kidney organoids

Human kidney organoids derived from our protocol have previously shown transcriptional association with trimester 1 human kidney, suggesting a relatively immature developmental phenotype (Takasato et al., 2016b). To analyse this at the single cell level, three independent kidney organoids were generated from the CRL1502 clone 32 female iPSC line as previously described (Takasato et al, 2016b) and profiled in parallel, generating a combined data set representing over 7000 cells, 6710 which passed quality control filtering based on the number of expressed genes and the expression of mitochondrial and housekeeping genes. An average of 2927 genes were expressed in each cell. This data has been submitted to NCBI’s Gene Expression Omnibus (Edgar et al, 2002) and is accessible through https://www.ncbi.nlm.nih.gov/geo/query/acc.cgi?acc=GSE108291. Unsupervised clustering of these cells using Seurat identified 16 populations with all cluster specific marker genes detailed in Supplementary Table 2 (Fig. 2A). Cells from replicate organoids were evenly distributed amongst clusters (Supplementary Figure 1C). Proposed cluster identities were assigned based upon expression of previously proposed markers (Fig. 2B). In addition, a Mann-Whitney test was used to examine the congruence between organoid and mouse developing kidney clusters from the two single cell datasets (Fig. 2C). This identified organoid populations representing early nephrons / podocytes (clusters 6 and 10), renal stroma (clusters 1, 2 and 5), vasculature (cluster 4), and a population expressing markers of various nephron segments and the ureteric epithelium (cluster 14). There were also lower confidence populations with similarity to nephron progenitors (cluster 13), early nephron and stroma (clusters 0, 8 and 9), stroma (cluster 7), and non-renal populations with similarity to neural cell types (clusters 11, 12 and 15) (Fig. 2A-C, Supplementary Figure 3).

**Figure 2.**
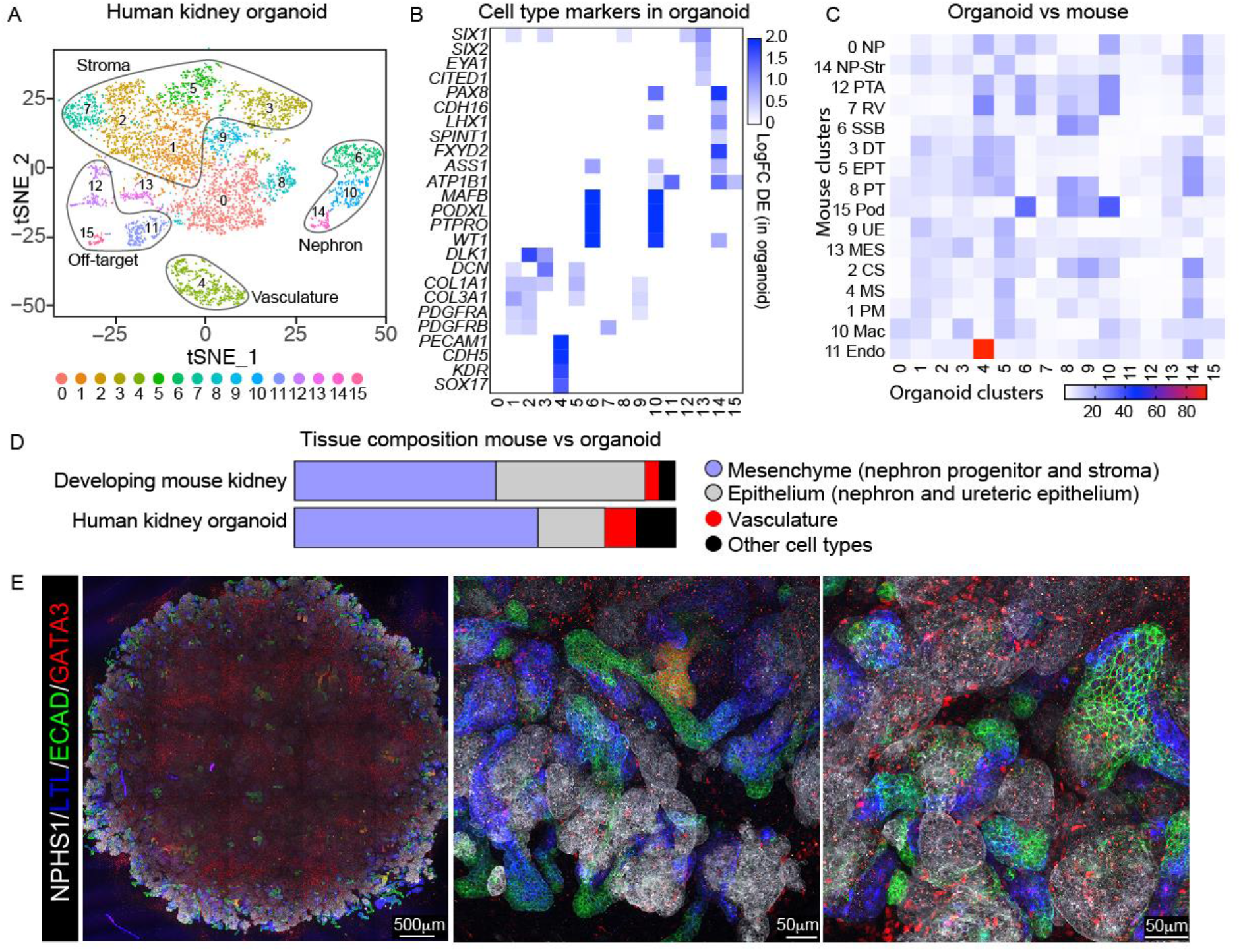
Single cell profiling of kidney organoids reveals both distinct renal and ‘off target’ cell types. **A**. tSNE plot idicating 16 distinct clusters including stroma (clusters 1,2,3,5,7), nephron (clusters 6,10,14), vasculature (cluster 4), nephron progenitor (cluster 13), unknown (cluster 0) and off target (clusters 11,12,15) cell types. Clusters 8 and 9 have some similarity to proliferating early nephron. **B**. Expression of key marker genes across cell clusters present within kidney organoids. **C**. Significance of overexpression of organiod marker genes across all kidney clusters using the non-parametric Mann-Whitney U test. Negative log10 p-values are plotted in the heat map. **D**. Analysis of the tissue composition of sequenced cells in the mouse kidney and organoids shown as the relative abundance of mesenchymal (nephron progenitor and stroma), epithelial (nephron and ureteric epithelium), vasculature and other cell types. **E**. Immunofluorescent staining of ‘sister’ organoids generated at the same time as those used for single cell transcriptional profiling. Antibodies used include NPHS1 (white; podocyte marker), LTL (blue; proximal tubule marker), ECAD (green; distal tubule marker) and GATA3 (red; marks collecting duct if ECAD^+^ and interstitial elements when ECAD^−^).

Recent single cell profiling of the adult mouse kidney identified the proximal tubule epithelium as the dominant cell type (Park et al., 2017). This agrees with standard histology of the postnatal organ, but is at odds with the developing tissue where the proximal tubules, and even the complete nephrons, represent a much smaller fraction of total cell due to an extensive interstitial compartment. As kidney organoids represent a model of the developing human kidney, we therefore compared the relative fraction of mesenchymal, nephron, endothelial and other cell types between the two datasets. This revealed a similar distribution of mesenchymal versus epithelial cell types in both structures (Fig. 2D). In order to correlate the cell types identified within the clusters with the specific organoids analysed, ‘sister’ organoids from the same differentiation were subjected to our standard immunofluorescence analysis. This identified the presence of significantly proximalised nephrons with a high proportion of podocytes (NPHS1^+^), although there was also clear evidence of presence of proximal tubule (LTL^+^, ECAD^+^), distal tubule (ECAD^+^GATA3^−^) and a small GATA3^+^, ECAD^+^ epithelial compartment we have previously defined as collecting duct (Fig. 2E). This is reflected in the relative size of the identifiable nephron clusters within the single cell data. There was also evidence of significant stromal GATA3 protein within the stromal compartment. This is also evident in the stroma of the developing mouse (Supplementary Figure 2).

### ‘Off target’ cell types within iPSC organoids

Several clusters identified within the kidney organoids did not correlate to native cell types in the developing mouse kidney (clusters 11, 12, 15). These populations had some similarity to neural progenitors (Supplementary Table 2) but lacked definitive markers to assign a specific cell type or state of differentiation based on currently available datasets. The presence of these ‘off-target’ cell types is not surprising considering the directed differentiation protocol commences with pluripotent stem cells. Embryonic derivatives of the epiblast include ectoderm, mesoderm, and endoderm-derived tissues. Neural cell types, regarded as an ectodermal derivative, are supported using a double inhibition of FGF and TGFβ signalling supported by low level fibroblast growth factor signalling (Lancaster and Knoblich, 2014), whereas renal cell types derive from the mesoderm (James and Schultheiss, 2003; Taguchi et al., 2014). As such, it is possible that ‘off target’ populations in our organoids represent cells that adopted a neural and perhaps even a neural crest identity during the early stages of differentiation and persisted in culture. Several other clusters within the organoids remain unlabelled, however an analysis of cell cycle suggests that cluster 8 and 9 represent cells in specific stages of the cell cycle (Supplementary Figure 1D). Results of using the Mann-Whitney U test to investigate the over expression of markers genes of organoid clusters in the kidney clusters showed some congruence between organoid clusters 8 and 9 and early nephron, which should represent a highly proliferative state. Finally, organoid cluster 0 had some similarity to stromal and mesenchymal cell types but no overlap with any kidney cell cluster.

### Subclustering of ureteric epithelium cells identifies known subpopulations and identifies developmental trajectories

We have previously described the formation of collecting duct / ureteric epithelium (PAX2^+^ECAD^+^GATA3^+^) within kidney organoids with the relative proportion of this epithelium dependent upon the duration of CHIR induction (Takasato et al., 2016b). In contrast, Taguchi et al propose that the collecting duct initially arises from a more anterior intermediate mesoderm and hence should not form simultaneously with the metanephric mesenchyme (Taguchi et al., 2014). The ureteric epithelium in the development mouse kidney has distinct zones of gene expression defining the tips, cortical, and medullary segments of this epithelium (Thiagarajan et al., 2011b). Cluster 10 expressed genes characteristic of ureteric epithelium, including *Wnt11, Ret, Hoxb7, Gata3*, and *Wnt9b*. Cells belonging to this cluster were re-clustered resulting in the identification of three sub-populations, with differential expression defining marker genes corresponding to tips, cortical, and medullary segments of the ureteric epithelium (Fig. 3A-C, Supplementary table 3) (Thiagarajan et al., 2011a). The cluster enriched for medullary gene markers contained genes expressed in the urothelium of the renal pelvis (Fig. 3C) (Thiagarajan et al., 2011a). Testing of Gene Ontology (GO), and Kyoto Encyclopaedia of Genes and Genomes (KEGG) annotations identified major signalling pathways active in the major sub populations. This indicated the activity of several pathways known to be involved in ureteric tip development such as WNT, retinoic acid, TGFβ, FGF, and YAP signalling (Fig. 3D, Supplementary Figure 4) (Reginensi et al., 2015; Yuri et al., 2017). Robust expression of *Gdnf* was detected in the nephron progenitor population and cortical stroma, in agreement with a recent report (Magella et al., 2017). Expression of GDNF receptors *Ret* and *Gfra1* were detected in the ureteric tip (Fig. 3FG, Supplementary Figure 2). This analysis also identified TGFB and PI3K-AKT pathways as active in the cortical collecting duct and phosphatidylinositol, PPAR, and Notch pathways as active in the medullary collecting duct and or urothelium (Fig. 3D). These represent candidate pathways for attempts to direct differentiation of mature collecting duct from progenitors of the ureteric epithelium. Pseudotime ordering of individual cells from kidney cluster 10 using Monocle (Trapnell et al., 2014) not only replicated the established developmental trajectory from tip progenitor to cortical then medullary collecting duct but identified additional states, and revealed cohorts of genes that change during this progression (Fig. 3E,F).

**Figure 3.**
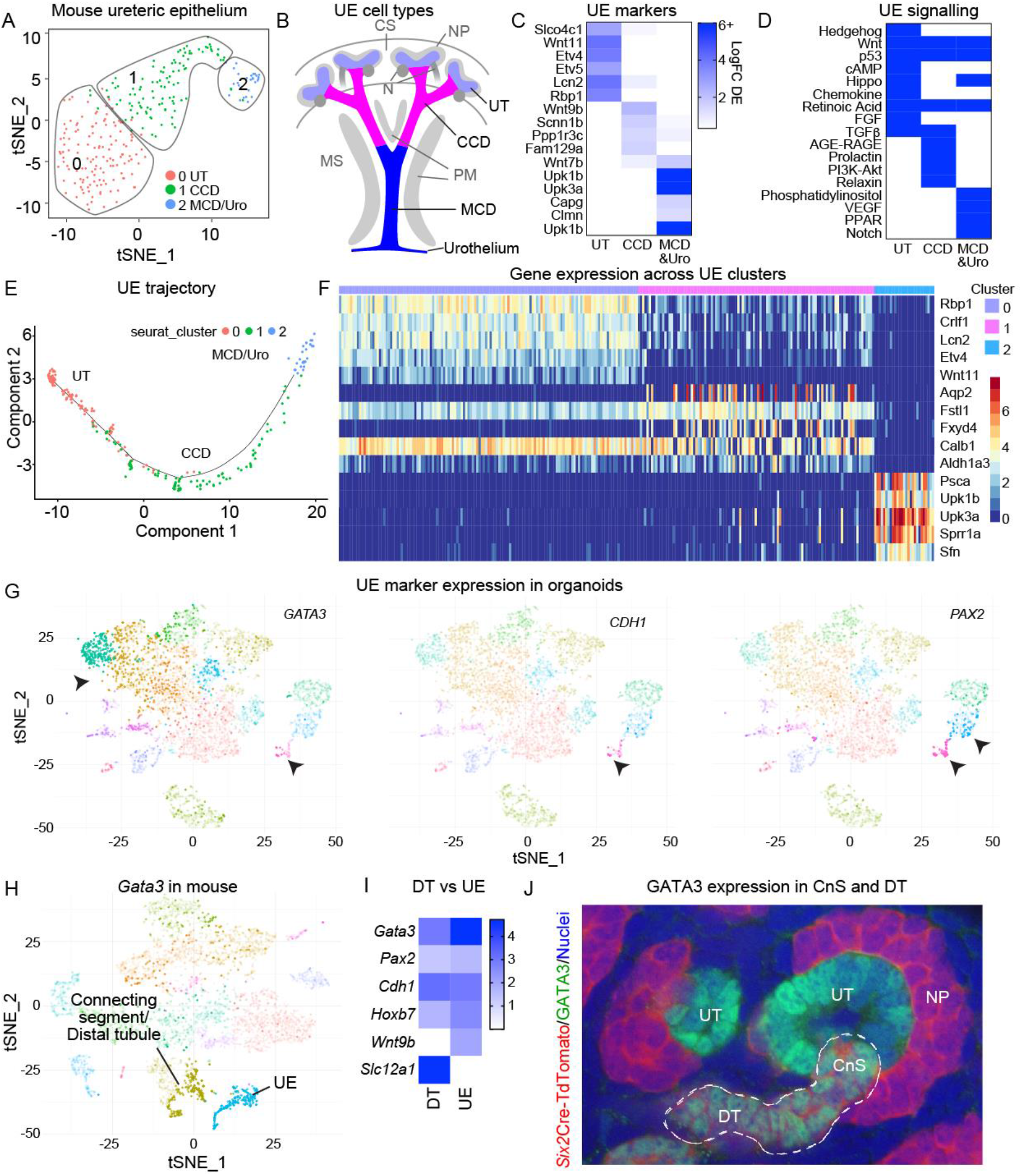
Subclustering of the collecting duct epithelium reveals marker overlap with distal tubule in mouse and organoid. **A**. Seurat reclustering of developing mouse ureteric epithelium (UE, Cluster 9) identifies three subclusters representing ureteric tip (UT, cluster 0), cortical collecting duct (CCD, cluster 1) and medullary collecting duct / urothelium (MCD/Uro, cluster 2). **B**. Diagram of the relative location of these three ureteric epithelial cell types with respect to surrounding stromal populations. CS, cortical stroma; NP, nephron progenitor; MS, medullary stroma; PM, pelvic mesenchyme. **C**. Expression of key marker genes of ureteric epithelium sub populations in UE subclusters. **D**. Identification of differential signalling pathway activity across these three UE populations. **E**. Monocle pseudotime trajectory of the three UE subclusters reflects a developmental origin of all clusters from the ureteric tip. **F**. Heat map of marker genes for subpopulations within the ureteric epithelium. Clusters represent UT (mauve, cluster 0), CCD (pink, cluster 1) and MCD (blue, cluster 2). **G**. Visualisation of key UE gene expression across tSNE plot of all profiled organoid cells revealing coexpression of CDH1, GATA3 and PAX2 in organoid Cluster 14. **H**. Visualisation of *Gata3* expression within the tSNE plot of developing mouse kidney shows expression in both distal tubule / connecting segment and ureteric epithelium. **I**. Relative expression of previously described markers of the collecting duct in distal tubule versus ureteric epithelium clusters within the developing mouse. This reveals promiscuous expression of most previously accepted collecting duct markers in distal tubule with only Wnt9b (UE) and Slc12a1 (DT) definitively identifying a single cell compartment. Scale indicates log fold change differential expression. **J**. High magnification image of an E15.5 Six2Cre-tdTomato mouse kidney showing co-staining of GATA3 (green) with *Six2*-derived nephron elements (red) specifically within the distal nephron / connecting segment (CnS).

### Reclassification of collecting duct within human kidney organoids

No cluster within the organoid data showed strong correlation with cluster 10 (UE) of the mouse developing kidney. Similarly, a specific analysis of the expression of key ureteric epithelium markers across the organoid t-SNE plot showed no evidence for a cell cluster with a clear ureteric epithelium identity (Fig. 3G). When examining organoids using immunofluorescence, we have previously defined the presence of ureteric epithelium on the basis of immunofluorescence for markers including PAX2, GATA3 and ECAD. A subset of cells within organoid cluster 14 showed clear expression of genes for these three proteins (*PAX2, GATA3* and *CDH1*), although there was also strong evidence for *GATA3* expression in stromal elements and *PAX2* expression in a wider set of epithelial cells (Fig. 3G). To determine whether the expression of these genes marked an epithelial population other than the ureteric epithelium in the developing mouse kidney, single gene analysis of the mouse kidney data was performed. This revealed clear evidence for the wider expression of many ureteric epithelium genes in mouse kidney cluster 3, which had been defined as Loop of Henle / distal tubule (Fig. 3H,I). Indeed, a direct comparison between the UE and DT clusters in mouse revealed highly congruent expression of a number of genes commonly regarded as marking UE (Fig. 3I). Few cluster specific markers were identified other than ureteric tip progenitors such as *Wnt11* and *Ret* (not shown), UE-expressed *Wnt9b* and DT-expressed *Slc12a1* (Fig. 3I).

To check that these results were not due to inappropriate clustering of ureteric epithelium cells in the mouse, we reexamined the presence of GATA3 protein within the distal nephrons segments (connecting segment / distal tubule) using a lineage reporting transgenic mouse line with a nephron progenitor-specific *Six2*-Cre (Kobayashi et al., 2008). As expected, all connecting segments were *Six2*-derived (nephron in lineage) but these structures were clearly GATA3+ protein (Fig. 3J). While expression of some ureteric epithelium markers in the connecting segment and distal tubule has been noted before (Georgas et al., 2008), the extent of the similarities has not previously been appreciated. This finding has implications for the identification of putative ureteric epithelium within kidney organoids. As a result, it is possible that the identity of the ECAD^+^GATA3^+^ epithelial segments in our kidney organoid represent distal tubule/connecting segment. It is curious to note that we have previously shown clear evidence for branching of this ECAD^+^GATA3^+^ epithelium within organoids, although nephrons do not branch. In addition, we never see evidence of formation of ureteric tips. This suggests that within organoids individual nephrons fuse to each other specifically at the distal end. This is a process potentially reminiscent of nephron arcading (al-Awqati and Goldberg, 1998), a process known to occur in the developing human kidney but has never been described in mouse. It should be noted that the organoids used for this specific experiment show proximal-biased nephrons. Hence it is possible that in some culture conditions a population of cells with ureteric epithelium identity is indeed present.

### Nephron lineages and relationships

Reclustering cells within the nephron lineage from the developing mouse kidney identified eight clusters as representing established early nephron states and mature nephron segments (Fig. 4A, Clusters 1-5,8,11,12) while accepted markers of the nephron progenitor state, including *Six2, Cited1*, and *Meox1*, defined five nephron progenitor clusters (clusters 0,6,7,9,10). All cells of the nephron arise from nephron progenitors via a mesenchyme to epithelial transition in response to tip-produced WNT9B (Carroll et al., 2005). The first morphological sign of this transition is clustering of progenitors into a pretubular aggregate (PTA), marked by expression of *Wnt4* and *Tmem100* (Rumballe et al., 2011). The PTA then differentiates into a polarised epithelial renal vesicle (RV), marked by *Ccnd1, Jag1*, and *Fgf8* (Fig. 4B) (Georgas et al., 2009). Patterning of the nephron tubule begins to emerge at the RV stage, with distinct gene expression patterns in proximal and distal segments (Georgas et al., 2009). Distal, medial, and proximal segments are evident in the s-shaped body (SSB) (Fig. 2B) (Georgas et al., 2008) with the proximal segment of the SSB body thought to give rise to the podocytes of the glomerulus (marked by *Mafb*) (Fig. 4B). The SSB matures into a capillary loop nephron, which contains precursors for all major nephron segments, including a connecting segment (*Calb1*), distal tubule (*Slc12a1*), loop of Henle (*Umod*), proximal tubule (*Slc22a6*), and podocyte-enriched glomerulus (Fig. 4B).

**Figure 4.**
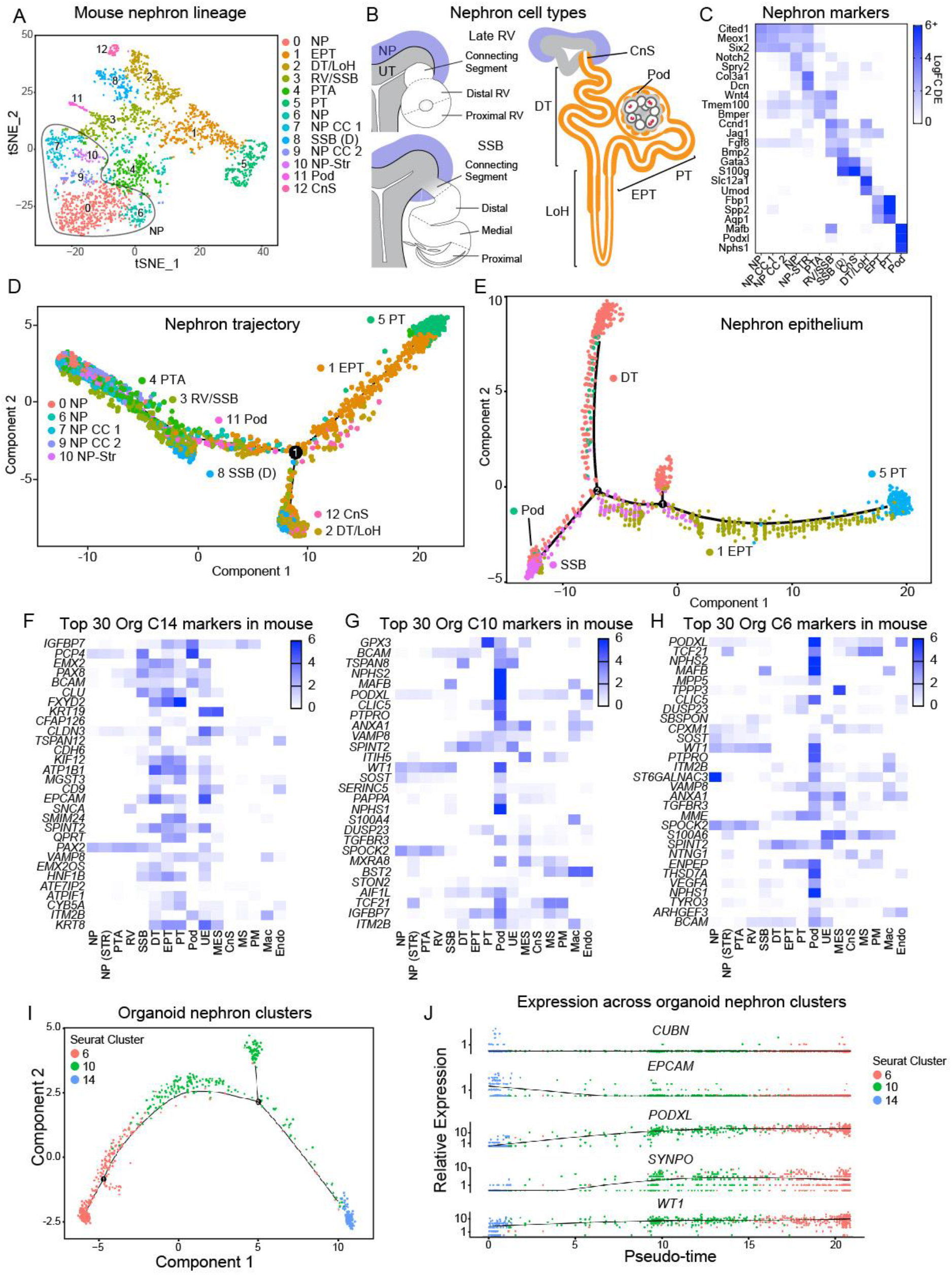
Comparisons between developing mouse kidney and human organoid nephrons reveals immature nephrons with a podocyte bias. **A**. tSNE plot of 13 selected cell clusters from developing mouse kidney representing cells of the nephron lineage. This includes 8 clusters representing distinct stages or segments of developing nephron (PTA, RV, SSB, EPT, Pod, PT, DT/LOH, CnS) including 5 clusters with nephron progenitor identity. **B**. Diagram of the stages of nephron maturation from the late RV through SSB and maturing nephron. Note the connecting segment that links the nascent nephron to the adjacent ureteric tip arises at late RV stage, by which time the distal and proximal RV already displays distinct gene expression (Georgas et al., 2009). By SSB, a medial domain of gene expression can be identified. A variety of genes have already been identified in the forming nephron to represent early PT versus PT (Thiagarajan et al., 2011b). **C**. Heatmap illustrating relative expression of key differentially expressed markers across these 13 nephron lineage cell clusters. **D**. Pseudotime analysis including of all cells in these clusters illustrates an anticipated transition from NP through PTA, RV/SSB, SSB distal (SSB (D). A clear branchpoint is observed between distal and proximal arms of nephron development. Of note, podocyte clusters are closer to RV/SSB than either proximal or distal tubule. **E**. Monocle pseudotime clustering using only the nephron cell clusters again places the majority of podocytes close to SSB and reveals a clear bifurcation between distal and proximal differentiation. In this instance, there is a sidebranch (1) which places a subset of DT cluster cells more closely related to SSB and early PT. **F-H**. Heatmaps displaying the expression of the top 30 differentially expressed genes for each of human iPSC-derived kidney organoid clusters 14, 10 and 6 (panels F, G and H respectively) relative to expression levels of these same markers across all 16 cell clusters in developing mouse kidney. This analysis suggests a more tubular identity for Cluster 14 with both Clusters 10 and 6 showing strongest similarity to podocyte. **I**. Monocle analysis of human iPSC-derived kidney organoid Clusters 6, 10 and 14 suggests a continuum of cell identity which places Cluster 14 as more distinct but also identifies a subpopulation within Cluster 10. **J**. Expression of tubule (*CUBN, EPCAM*) and podocyte (*PODXL, SYNPO, WT1*) genes across these three nephron cell clusters within organoids again suggests a more tubular identity for Cluster 14 with strong similarity in profile between Clusters 10 and 6.

Cells representing all major nephron segments were present in the mouse single cell data (Fig. 4C), with RV and SSB markers overlapping in cluster 3, and cluster 8 aligning most closely with the distal (DT) and connecting segments (CnS) of the SSB body. Clusters representing distal tubule and loop of Henle (*Slc12a1, Umod*), early proximal tubule and proximal tubule (*Fbp1, Spp2*), and podocytes (*Mafb, Podxl*) were also identified (Fig. 4C). While classification of cell types relied upon established marker genes, the accompanying lists of differentially expressed genes between clusters offers insight into novel markers (Supplementary Table 4).

In order to interrogate nephron formation further we performed pseudotime analysis using Monocle. This identified three main states within the nephron lineage (Fig. 4D). An initial state combined nephron progenitors, early nephron states up to SSB and podocytes. The trajectory subsequently forked into two arms representing the connecting segment and distal tubule on one arm, and the proximal tubule on the other (Fig. 4D). Assessing just clusters from the nephron epithelium placed podocytes alongside SSB cells, and again divided distal tubule from proximal tubule (Fig. 4E). This trajectory is consistent with our current understanding of early nephron patterning and podocyte specification (Georgas et al., 2009; Georgas et al., 2008).

Within kidney organoids, three clusters represented nephron epithelium (clusters 6, 10 and 14). Cluster 14 expressed markers of the nephron and/or ureteric epithelium, with co-expression of markers enriched in the SSB (*LHX1*), distal tubule (*SPINT2, CDH16*), proximal tubule (*HNF1B, ASS1*), and ureteric epithelium (*EPCAM, KRT19*) of the developing mouse kidney (Fig. 4F). Nephrons within human kidney organoids represent immature nephrons, up to the capillary loop stage (Takasato et al., 2016b). During early nephron formation, markers of distinct mature nephron segments can be co-expressed (Georgas et al., 2009; Thiagarajan et al., 2011b). Nephron patterning in the organoids profiled in this particular study was skewed towards a proximal fate (Fig. 2E), with a higher proportion of podocyte and proximal tubule specification, and lower distal nephron representation than organoids previously generated by this protocol (Takasato et al., 2014; Takasato et al., 2016b). This discrepancy may reflect batch effects in media composition or growth factor efficacy. Clusters 6 and 10 both showed strong expression of podocyte markers including *MAFB, PODXL, PTPRO, NPHS1, NPHS2* and strong similarity with the podocyte cluster in mouse (Fig. 4G,H; Fig. 2BC; Supplementary Figure 3). We would note, however, than many podocyte transcription factors, including *WT1, MAFB* and *NPHS1*, are expressed from very early in nephron formation hence these two clusters may represent early and later proximal nephron. Monocle pseudotime analysis of organoid nephron clusters (6, 10, and 14) (Fig. 4I) indicated a trajectory from cluster 14 through 10 to 6 with a sidebranch from cluster 10. As podocytes appeared to map as an early sidebranch from proximal and distal tubule clusters in mouse, we examined whether this sidebranch of cluster 10 cells showed stronger podocyte gene expression. This was not the case. However, tubular markers (*CUBN, EPCAM, CDH16*) were highest in cluster 14 and gave way to podocyte markers (*PODXL, SYNPO, WT1*) in clusters 10 and 6 (Fig. 4J).

### Identification of a novel nephron progenitor state in mouse kidney and human kidney organoids

Nephrons form from a self-renewing mesenchymal population marked by expression of *Six2* and *Cited1* (Boyle et al., 2008; Kobayashi et al., 2008). Currently, the nephron progenitor population is thought to be divided into two positionally distinct subdomains, a *Six2+Cited1+* uncommitted state and a *Six2+Cited1^−^* state sensitised to commitment to pretubular aggregate (Brown et al., 2015). However, previous timelapse imaging of kidney morphogenesis has revealed substantial cell movement (Combes et al., 2016) and variation in cell cycle length (Short et al., 2014) within the nephron progenitor population, suggesting this cell population may be more heterogeneous than previously thought. As noted above, five nephron progenitor populations were identified within the mouse data. Two of these (cluster 7 - DNA replication and cluster 9 - mitosis) appeared to be driven by cell cycle genes, despite this factor being regressed from the dataset prior to clustering. Both of these clusters displayed a more committed phenotype with detectable, albeit low, expression of *Wnt4* and *Tmem100*. This could suggest that these clusters are primed for commitment and cell division may be a requirement for this commitment process. The three remaining nephron progenitor clusters all expressed canonical nephron progenitor genes as well as cluster specific differentially expressed markers (Fig. 5A). We define these here as 1) nephron progenitor ‘ground-state’ (cluster 0) with little to no expression of *Wnt4* and *Tmem100*, 2) nephron progenitors (cluster 6) with slightly lower levels of *Six2* and *Cited1* and coexpression of *Notch2* and *Sprouty2* and 3) a novel nephron progenitor population (cluster 10) with modest expression of commitment markers and clear stromal characteristics including expression of *Pdgfra* and *Col3a1* (Fig. 5AB). We define Cluster 10 as NP-STR and show a clear intermediate transcriptional profile for this cluster that sits between known NP and stromal populations (Fig. 5B). With respect to cluster 6, *Sprouty2* is a negative regulator of FGF signalling, with FGF signalling associated with nephron progenitor maintenance (Walker et al., 2016). Notch signalling has recently been shown to regulate early commitment to nephron formation (Chung et al., 2016; Chung et al., 2017). Hence, cluster 6 may represent a transitional state between nephron progenitor and pretubular aggregate. It has been presumed based on IF and section *in situ* hybridisation, that a portion of the nephron progenitor population was SIX2^+^CITED1^−^, with these cells more committed to nephron formation. At the single cell level, no *Six2^+^Cited1^−^* progenitor state was observed although this may be due to regulatory mechanisms acting after transcription or translation.

**Figure 5.**
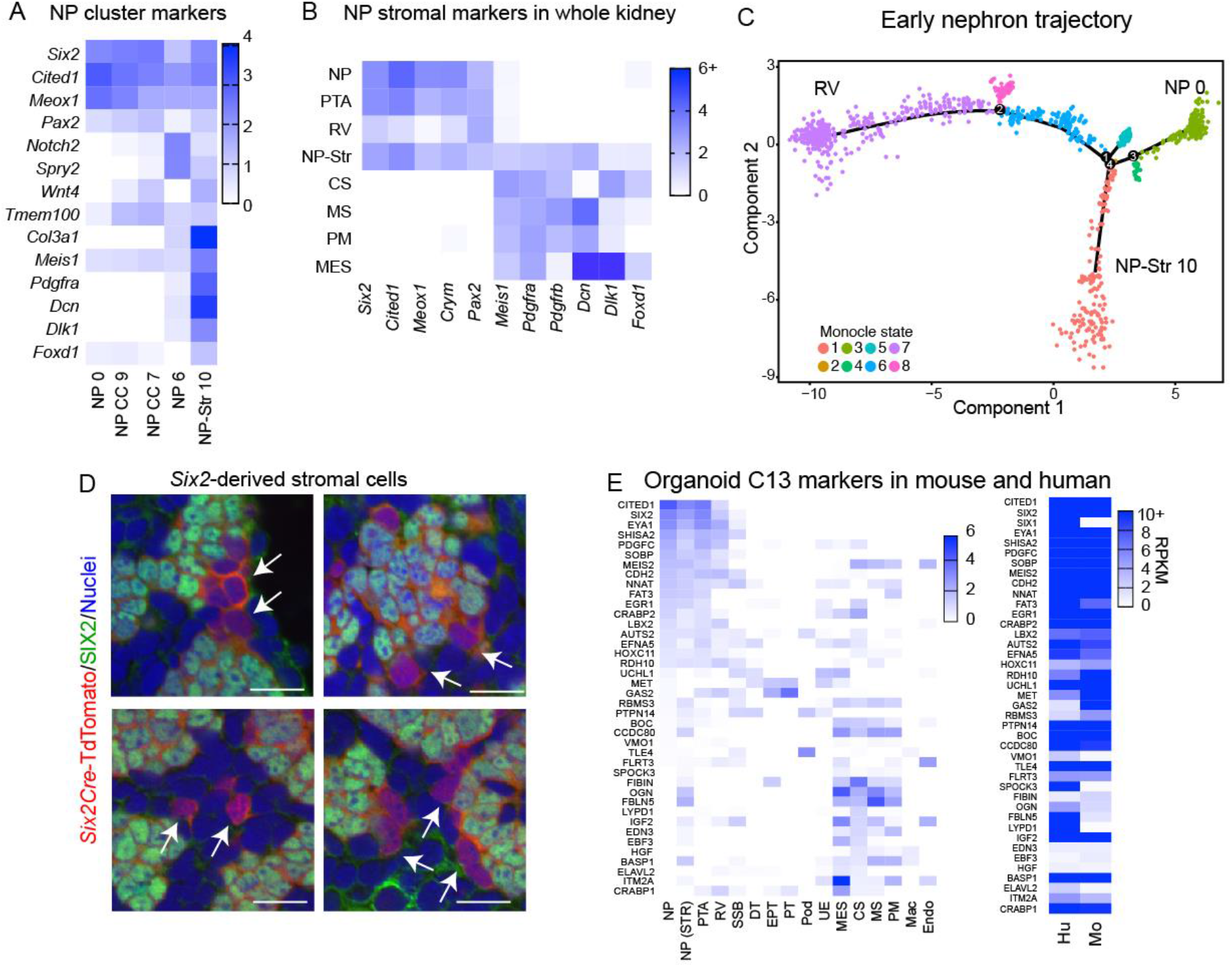
Transcriptional evidence for substantial nephron progenitor heterogeneity and novel NP-stromal subclusters. **A**. Expression of key markers and distinguishing genes between nephron progenitor clusters from the nephron lineage clustering in Fig. 4*A*. **B**. Expression of nephron progenitor and stromal markers in early nephron and stromal clusters from the whole kidney demonstrates a nephron progenitor population with a clear stromal signature. Scale represents log fold change differential expression. **C**. Monocle analysis of early nephron lineage clusters identifies a distinct developmental trajectory for nephron progenitor-stromal cells, distinct from the expected NP-RV trajectory taken by most cells. **D**. Lineage tracing from the *Six2* locus identifies *Six2*-derived cells (red), negative for SIX2 protein (green), in the stroma between nephron progenitor niches. **E**. Left-expression of the top 40 markers of organoid cluster 13 in the developing mouse kidney, sorted by expression in NP and CS populations. Scale is log fold change differential expression in mouse. Right-expression of the same markers in bulk RNA-Seq data from ITG8A^+^ nephron progenitor-enriched cells from human fetal kidneys (Hu), and bulk RNA-Seq data from CITED1^+^ nephron progenitors from the developing mouse kidney. Scale indicates reads per kilobase of transcript per million mapped reads from 0-10 and above.

We next investigated how the new nephron progenitor states, clusters 6 and 10 (NP-STR), fit with the established developmental trajectory from nephron progenitor to renal vesicle using pseudotime analysis. Assessing cells within early nephron clusters (NP to RV, clusters 0,4,6,7,9,10; Fig.5C) reproduced the expected developmental trajectory from nephron progenitor to pretubular aggregate and renal vesicle, with additional subpopulations on minor side branches corresponding to cell cycle state (Fig. 5C). The NP-STR state diverged from this main trajectory immediately after the nephron progenitor ground state.

Nephron progenitors and stromal progenitors arise from the same lineage before the onset of nephron formation (Brunskill et al., 2014; Mugford et al., 2008) but are not thought to cross between lineages under normal conditions (Kobayashi et al., 2014; Naiman et al., 2017). The observation of a nephron progenitor state with stromal marker expression could represent cells transitioning between the nephron and stromal lineages. To investigate this, we returned to *in vivo* lineage tracing experiments using nephron progenitor expressed Six2-Cre. Here we observed the rare but consistent presence of Six2-lineage labelled cells within the renal stroma (Fig. 5D). While these cells were negative for *Six2* protein, they clearly have arisen from the *Six2*^+^ nephron progenitor population, supporting the existence of a transitional cell type. Coexpression of nephron progenitor and stromal markers has been noted in rare cells at earlier developmental times (Brunskill et al., 2014; Magella et al., 2017). The persistence of a nephron progenitor state with a stromal transcriptional signature at this late developmental stage could indicate that nephron progenitors are susceptible to external signals that promote stromal fate. Alternatively, nephron progenitors may default to a stromal fate if they do not receive sufficient signals from the nephron progenitor niche. *Pax2* has recently been proposed to repress stromal identity in nephron progenitors as these cells transdifferentiate to stroma in the absence of this gene (Naiman et al., 2017). The nephron progenitor-stroma cluster co-expresses *Pax2* and markers of stromal fate, indicating that other genes are involved in maintaining nephron progenitor identity.

While our current kidney organoid protocol shows evidence of nephron formation and segmentation, the peak of SIX2 expression was low and early during the differentiation protocol (day 10) and was followed by a significant decline in all nephron progenitor markers with time (Takasato et al, 2015). Antibody staining did not reveal strong evidence for a strong nephron progenitor domain, suggesting that nephron formation was initiated and then proceeded without the maintenance of the progenitor population. A comparison of the NP markers from the mouse kidney to clusters within the kidney organoids suggested that Cluster 13 showed the strongest similarity to NP with the expression of *CITED1, MEOX1, SIX2, SIX1, EYA1*, and *CDH2* (Fig.2B; Fig.5). While GO analysis showed an association to neural development, co-expression analysis using ToppFun (https://toppgene.cchmc.org/enrichment.jsp) identified a nephron progenitor signature. Although *SIX1* and *EYA1* play important roles in specifying neurons (Zou et al., 2004) and nephron progenitors (Xu et al., 1999; Xu et al., 2003), this organoid cluster lacked definitive neural markers such as *NEUROG1*. 73% of markers differentially expressed with a log-fold change greater than 0.3 within this cluster were expressed in nephron-progenitor enriched ITG8A^+^ cells from human fetal kidneys (Fig. 5E) (O’Brien et al., 2016). Interestingly, this putative nephron progenitor population also had stromal characteristics, expressing several genes restricted to stromal populations in the developing mouse kidney such as *ITM2A, OGN, IGF2, CRABP1*, but lacking *PDGFRA* and *MEIS1* (Fig.5E). As such, this population showed similarity to the novel NP-STR state (and by inference the stromal clusters) in the developing mouse kidney (Fig. 5E). This would support a model in which nephron progenitors represent a plastic cellular identity with appropriate spatial signalling from an adjacent ureteric tip suppressing expression of stromal markers.

### Lessons on stromal subpopulations within developing kidney and organoids

Relatively little is known about the molecular characteristics of stromal populations within the developing kidney. To explore stromal heterogeneity in the developing mouse kidney further, we reclustered cells from the stromal lineage alone, resulting in seven clusters (Fig. 6A). New clusters were mapped back to those identified in the whole kidney to assist in cluster identification (Supplementary Figure 5A). Stromal cluster 0 is likely cortical stroma (*Foxd1*^+^) and the expression signature of related cluster 4 is likely driven by cell cycle features. Stromal clusters 2 and 3 likely represent sub populations of pelvic mesenchyme (*Wnt11*+), 1 and 5 are related to medullary stroma, and stromal cluster 6 maps directly kidney cluster 13, which may represent mesangial cells (Fig.6B, Supplementary Figure 5A). The lists of differentially expressed genes in each population resulting from this analysis will facilitate the characterisation of stromal populations in the developing kidney (Supplementary Table 5). Monocle analysis of these stromal clusters identifies a progression from cortical stroma through intermediate populations to a split between pelvic mesenchyme, medullary stroma and a putative mesangial population (Supplementary Figure 5B).

**Figure 6.**
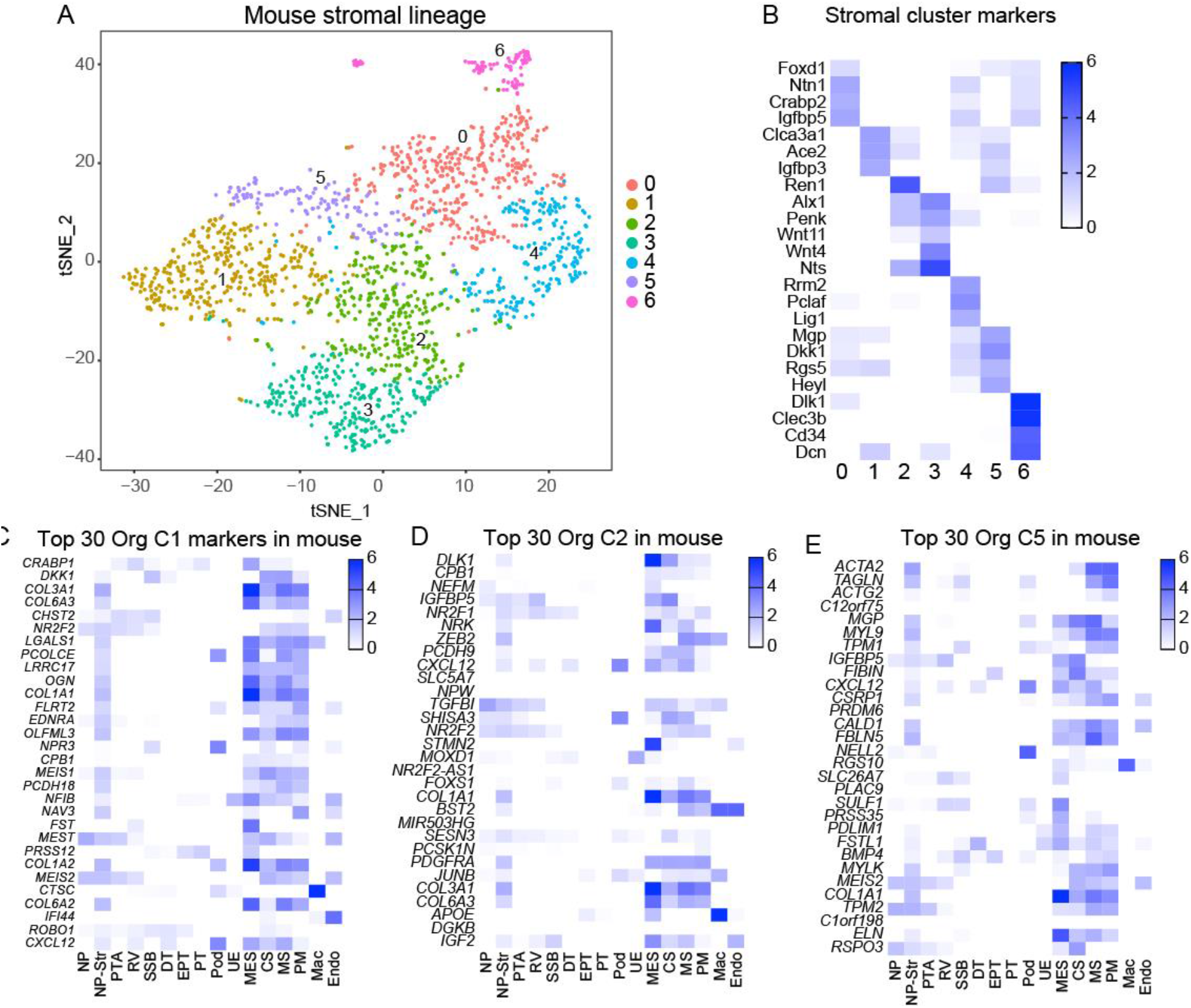
Analysis of stromal clusters within developing mouse kidney and human iPSC-derived kidney organoids. **A**. Reclustering of all cells from the stromal lineage from the developing mouse kidney resulted in seven clusters. **B**. Marker genes differentially expressed between stromal clusters. **C-E**. Heatmaps displaying the expression of the top 30 differentially expressed genes for each of human iPSC-derived kidney organoid clusters 1, 2 and 5 (panels C, D and E respectively) relative to expression levels of these same markers across all 16 cell clusters in developing mouse kidney. This analysis shows expression of almost all of organoid cluster 1 markers in the stroma of the developing mouse kidney (Mes, CS, MS, PM clusters), and similar expression of several markers from organoid clusters 2 and 5.

Multiple organoid populations were identified that expressed markers restricted to stromal cell types in the developing mouse kidney with this match being strongest in organoid clusters 1, 2, and 5 (Fig. 6C-E). Common markers included *COL3A1, PDGFRA, PDGFRB* and *MEIS1*. Other genes differed between these clusters including *DKK1* (expressed in organoid cluster 1) *DLK1* (expressed in organoid cluster 2) and *DCN* (expressed in organoid cluster 3). In the developing mouse kidney, *Dkk1* is expressed in cortical and medullary stroma, *Dlk1* is expressed in the cortical stroma, and *Dcn* is expressed in the medullary stroma and pelvic mesenchyme. While these differences in marker gene expression suggest similarity to distinct stromal populations in mouse, this was not fully supported by broader analysis of the expression profiles.

### Mechanisms regulating nephron formation and maturation

Transcriptional regulation is a critical mechanism for determining and maintaining cell fate during development. Segment specific cohorts of transcriptional regulators may be informative for direct reprogramming as previously reported for NP and proximal tubule respectively (Hendry et al., 2013; Kaminski et al., 2016). The top differentially expressed transcription factors within each mouse nephron lineage cluster, including nephron progenitors, were identified (Supplementary Figure 4, Supplementary Table 4), highlighting cell type-specific transcription factors such as *Six2* (nephron progenitor and early nephron), *Mafb* (renal vesicle, podocytes), and *Hnf4a* (proximal tubule) (Kaminski et al., 2016; Thiagarajan et al., 2011a). Signalling pathways identified as active within the nephron progenitor cluster include several pathways shown to regulate nephron progenitor fate *in vivo* such as PI3K-AKT, WNT, Hippo and MAPK signalling (Brown et al., 2015; Das et al., 2013; Karner et al., 2011; Lindstrom et al., 2015; McNeill and Reginensi, 2017). Likewise, signalling pathways capable of triggering nephron formation from the progenitor state such as Notch and TGF-β signalling (Brown et al., 2015; Chung et al., 2017) were identified in early nephron cell types (Supplementary Figure 4). In addition to these known players, other developmentally significant signalling pathways such as Hedgehog and JAK-STAT are implicated by this analysis and could be used to improve recently developed methods to culture isolated nephron progenitors (Brown et al., 2015; Li et al., 2016; Tanigawa et al., 2016). While nephron progenitor regulation has been intensely studied, very little is known about the signals controlling the maturation of nephron segments. This analysis identifies cAMP, cGMP-PKG, and insulin signalling as candidates to promote formation of distal tubule and early proximal tubule, and PPAR signalling (facilitates non-canonical retinoic acid signalling) as an additional candidate for proximal tubule maturation. Specific information about ligands and receptors is available (Supplementary Table 4). This data provides candidate signalling pathways that could be used to produce specific states of nephron maturation from primary nephron progenitor cells and in human kidney organoids.

## Discussion

The developing kidney represents an invaluable tool with which to understand the formation and maturation of each renal cell type. The single cell data presented here offers a unique opportunity to comprehensively understand the mechanisms of progenitor maintenance and differentiation in the stroma, the ureteric epithelium, and the nephron lineages. Dynamic changes in transcription factor and signalling pathway activity from progenitor to mature cell type are revealed in all three lineages and therefore provide a roadmap of the signals that regulate progenitor maintenance and differentiation *in vivo*. This represents a major advance and provides a unique opportunity to utilize this novel insight into the molecular regulation of renal development to refine strategies to control the directed differentiation of renal cell types. This information can be used to refine attempts to maintain progenitor populations and direct differentiation of both primary and pluripotent cells.

This is not the first application of single cell profiling to the developing mouse kidney. Indeed, previous analyses of E12.5 mouse kidney using the Fluidigm platform identified evidence for multi-lineage priming of nephron progenitor fate and early plasticity between nephron and stromal lineages (Brunskill et al., 2014). The mechanisms of nephron progenitor ageing were also interrogated in this way (Chen et al., 2015). More recent studies applying the DropSeq approach examined far larger numbers of individual cells in younger (14.5dpc) kidneys showing that expression of *Gdnf*, a key signalling molecule that drives ureteric bud branching, is expressed in a broader population of cells than previously thought (Magella et al., 2017). Another study warned of non-specific gene expression changes in response generating single cell suspensions using proteases active at 37 degrees Celsius (Adam et al., 2017). Our analyses using the 10x Chromium platform align with these previous reports but identify a larger number of cell clusters. This has particularly facilitated a more granular analysis of cell types within the ureteric epithelium and the nephron progenitor population. Indeed, we report here the identification of a clear nephron progenitor-stromal sub-cluster predicted by pseudotime analysis to contribute to both the cap mesenchyme and nephron formation.

More importantly, this work also addresses a timely question of how faithfully stem cell-derived tissues represent native cell types. As for all iPSC-derived organoid protocols described to date, the kidney organoids most likely represent models of human trimester 1 and 2, hence the developing kidney is a more accurate comparator than adult kidney, either mouse or human. By comparing differentially expressed gene profiles for each cellular subcompartment, we have been able to identify and classify as renal 8 of the 16 clusters within the organoids profiled. By comparing kidney organoid cell types with the developing mouse kidney, we reinforce findings to date suggesting that kidney organoids represent an immature nephron state, with podocytes and endothelial cells most closely aligning to equivalent mouse cell types. Stromal and nephron lineages are present but subpopulations are not as clearly defined, perhaps due to a lack of appropriate spatial patterning resulting from three dimensional growth and cellular interactions that occur *in vivo*. While these limitations have been acknowledged before, we note that several components of the developing kidney are represented in this model. Importantly, this study has redefined the distal segment of the nephron present in this protocol and characterised the ‘off target’ endpoints present. Overall, the data obtained here will serve as a base from which to optimise the differentiation of kidney organoids to build a better model of the developing human kidney. Based on the detailed analysis of the transcription factors expressed and the signalling pathways active in specific cell types during mouse development, we also have a roadmap with which to further mature the organ so as to develop a more accurate model of the postnatal organ.

## Methods

### Mouse Strains and Embryo Staging

In mouse experiments, noon of the day on which the mating plug was observed was designated embryonic day (E) 0.5. C57Bl/6 mice were used for the E18.5 embryonic kidney analysis. Tg(Six2-EGFP/cre)1Amc mice were mated to Gt(ROSA)26Sor^tm9(CAG-tdTomato)Hze^ homozygotes to generate E18.5 embryonic kidneys with tdTomato labelling in the nephron lineage. All animal experiments in this study were assessed and approved by the Murdoch Children’s Research Institute Animal Ethics Committees and were conducted under applicable Australian laws governing the care and use of animals for scientific purposes.

### Immunofluorescence and Microscopy

E18.5 embryonic kidneys were fixed in 4% PFA for 20 minutes, washed in PBS and cleared using the PACT method (Yang et al., 2014) to preserve tdTomato fluorescence. Cleared samples were stained using rabbit anti-Six2 (Proteintech, 11562-1-AP) or Goat anti-Gata3 (R&D Systems, AF2605) and Alexa Fluor 488 labelled secondary antibodies (Thermo Fisher). Samples were blocked in PBST (PBS + 0.1% Triton-X) with 10% normal donkey serum and incubated at room temperature with each antibody solution for at least 48 hours followed by washing for 24 hours in PBST. Nuclei were stained using Draq5 (Abcam). Samples were mounted in RIMS (88% Histodenz) and imaged using an Andor Dragonfly spinning disk system with a 40*um* pinhole disk and Nikon 1.15NA 40x water-immersion objective. Images were processed in Fiji (Schindelin et al., 2012).

### Organoid differentiation

Kidney organoids were made according to our published protocol (Takasato et al., 2016a) from human induced pluripotent stem cell line CRL1502 (Briggs et al., 2013). Three organoid samples were differentiated to day 25 (7 days of monolayer culture +18 days as a 3D aggregate).

### Single cell sample prep and sequencing

Mouse kidneys were dissected into ice cold PBS then digested over 15 minutes at 37C in Accutase (#A1110501 Life technologies), with manual dissociation via pipetting through a P1000 tip every 5 minutes. Following dissociation, cells were passed through a 30 micron filter and stored on ice in 50% PBS, 50% DMEM with 5% FCS. Human kidney organoids were dissociated via a similar protocol using Trypsin (#25300054, Life Technologies) instead of Accutase and stored on ice in APEL media. Three pairs of 18.5 dpc mouse kidneys and were run in parallel on a chromium 10x Single Cell Chip (10x Genomics), as were 3 independent organoids in a separate run. Libraries were prepared using Chromium Single Cell Library kit V2 (10x genomics), and sequenced on an Illumina HiSeq using 100bp paired-end sequencing.

### Data submission information

The data discussed in this publication have been deposited in NCBI’s Gene Expression Omnibus (Edgar *et al*., 2002) and are accessible through GEO Series accession number GSE108291 (https://www.ncbi.nlm.nih.gov/geo/query/acc.cgi?acc=GSE108291).

### Single cell data analysis

#### Mouse

Raw sequencing data was processed using Cell Ranger (v1.3.1, 10x Genomics) to produce gene-level counts for each cell in each sample, which were aggregated to form a single matrix of raw counts for 6752 cells. All subsequent analysis was performed in the R statistical programming language. Cells with greater than 95 per cent of genes with zero assigned reads were removed. Genes with counts in less than 50 cells, mitochondrial and ribosomal genes, and genes without annotations were also filtered out. This approach collectively removed the majority of blood cells from the data set, however signatures of remaining blood cells can be seen in the final data. The final dataset used for analysis consisted of 5639 cells and 13116 genes. The Seurat package (v2.0.1) (Macosko et al., 2015; Satija et al., 2015) was used to normalise data, regressing out factors related to biological replicate and cell cycle. For clustering, 1962 highly variable genes were selected and the first 30 principal components based on those genes used to build a graph, which was segmented with a resolution of 0.8. We obtained lists of differentially expressed genes for each cluster by testing for genes that had a log foldchange greater than one between cells in one cluster compared with all other cells using the glmTreat method in the edgeR package (Robinson et al., 2010). GO and KEGG analysis was performed with limma (Ritchie et al., 2015). Trajectory analysis of the various lineages was performed using Monocle (v2.4.0) (Trapnell et al., 2014).

#### Human kidney organoid

Cell Ranger was used to process and aggregate raw data from each of the samples returning a count matrix for 7004 cells, which was then read into R. The scater package (v1.4.0) (McCarthy et al., 2017) was used to produce quality control plots. Cells that expressed more than 8000 genes (possibly representing doublets), had more than 9 per cent of reads assigned to mitochondrial genes or showed low expression of ACTB or GAPDH were removed. Genes that were expressed in less than two cells or had less than two counts across the whole dataset were filtered out as were mitochondrial genes, ribosomal genes, those associated with the cell cycle and those without HGNC symbols. The final dataset had 6710 cells and 13832 genes. Factors associated with the total counts in each cell and the percentage of counts assigned to mitochondrial genes were regressed out of the dataset and 1612 highly variable genes were selected for clustering. The first 20 principal components based on these genes were selected to build a graph, which was clustered with a resolution of 0.8. Seurat’s likelihood-ratio test was used to determine if genes were differentially expressed between each cluster and all other cells, only testing those genes expressed in at least 25 per cent of cells in a cluster with a log fold change greater than 0.25. Monocle was used to perform trajectory analysis.

## Author contributions

P.E. performed all differentiations, single cell isolation and immunofluorescence evaluation of organoids. B.P. and L.Z. performed the single cell mapping, normalisation, differential expression and trajectory analysis on the mouse and human organoid data respectively, under the supervision of A.O. K.T.L performed additional data analysis and experimental validation. A.N.C performed single cell experiments and was primarily responsible for interpretation of the data. A.N.C. and M.H.L conceived and designed the experiments and wrote the manuscript. All authors contributed intellectually to the project and manuscript revisions.

## Acknowledgements

This work was supported by the Australian Research Council (DE150100652), National Institutes of Health Rebuilding a Kidney consortium (DK107344) and seed funding from the Murdoch Children’s Research Institute and the University of Melbourne. Single cell sequencing was performed at the Australian Genome Research Facility Genomics Innovation Hub with the assistance of J. Jabbari and A. Seidi. Microscopy was performed at the Murdoch Children’s Research Institute. A.N.C. holds a Discovery Early Career Researcher Award from the Australian Research Council. M.H.L. is a Senior Principal Research Fellow of the NHMRC. A.O. is a Career Development Fellow of the NHMRC. L.Z. is supported by an Australian Government Research Training Program (RTP) Scholarship. MCRI is supported by the Victorian Government’s Operational Infrastructure Support Program.

## Supplementary Data

**Supplementary Table 1. Interactive spreadsheet containing lists of differentially expressed genes in all 16 clusters within the E18.5 developing mouse kidney. Use the “lookup” tab to access the interactive sheet and input an official gene symbol in the left column to retrieve differential expression results for that gene across all clusters**.

**Supplementary Table 2. List of marker genes identifying 16 cell clusters within human kidney organoids**.

**Supplementary Table 3. Interactive spreadsheet containing lists of differentially expressed genes identifying 3 ureteric epithelium subclusters within the E18.5 developing mouse kidney**.

**Supplementary Table 4. Interactive spreadsheet containing lists of differentially expressed genes identifying 8 nephron lineage and 5 nephron progenitor subclusters within the E18.5 developing mouse kidney**.

**Supplementary Table 5. Interactive spreadsheet containing lists of differentially expressed genes identifying 6 stromal subclusters within the E18.5 developing mouse kidney**.

**Supplementary Figure 1.**
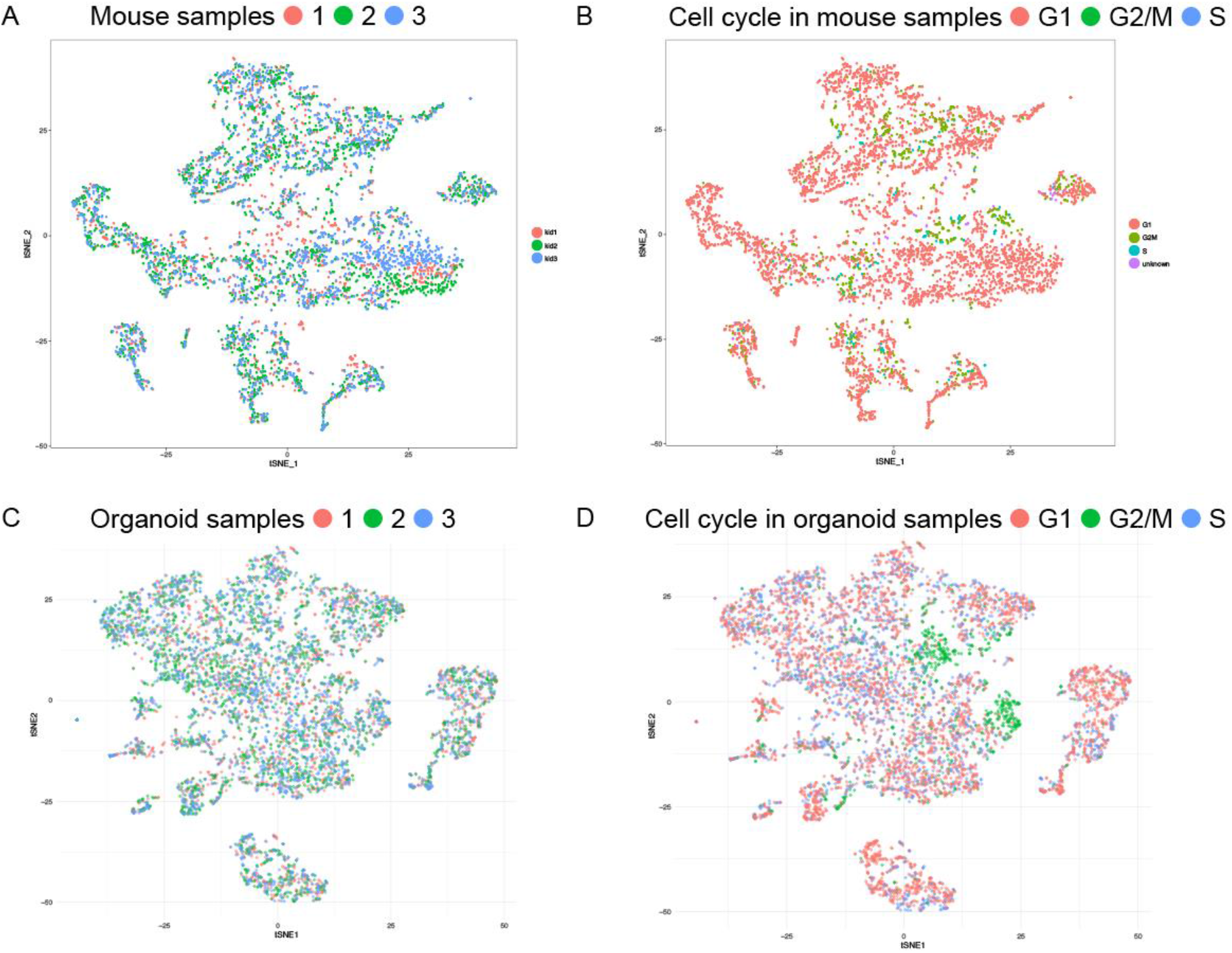
Visualisation of all single cell data by sample and cell cycle. **A**. tSNE plot of all cells from mouse developing kidney identified by individual mouse samples. This shows an even distribution of cell types present within all samples. **B**. tSNE plot of all cells from mouse developing kidney identified by stage of cell cycle (G1, G2/M, S). **C**. tSNE plot of all cells from human iPSC-derived kidney organoids identified by individual organoid. This shows an even distribution of cell types present within all samples speaking to the reproducibility between organoids. **D**. tSNE plot of all cells from human iPSC-derived kidney organoids identified by stage of cell cycle (G1, G2/M, S). This suggests that the identification of cluster 8 and 9 is based on cell cycle.

**Supplementary Figure 2.**
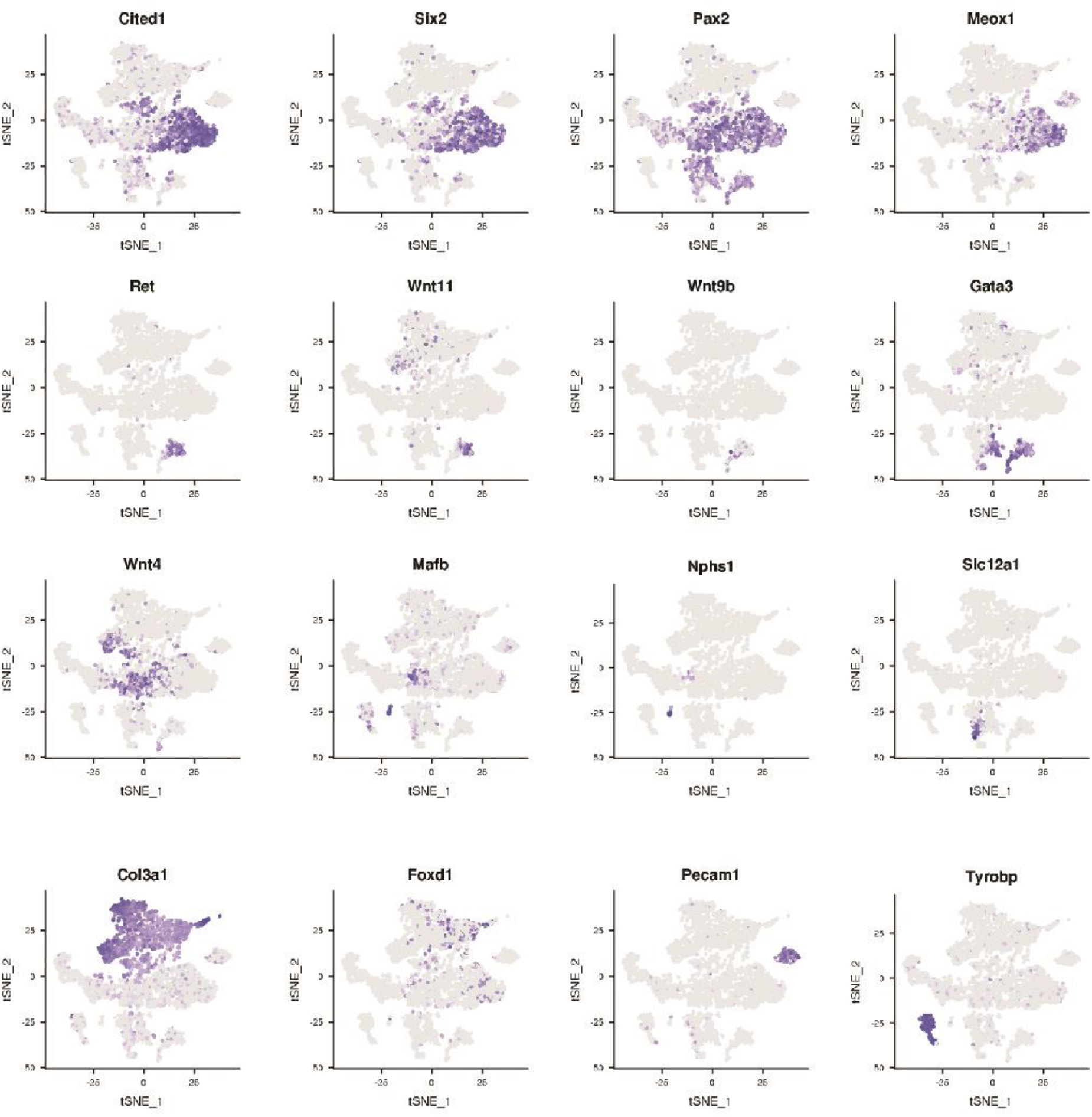
Marker expression across cells in the developing mouse kidney. tSNE plot of all cells from E18.5 developing mouse kidney showing the location of cells expressing key marker genes, including markers of nephron progenitor (*Cited1, Six2, Pax2, Meox1*), ureteric epithelium (*Ret, Wnt11, Wnt9b, Gata3*), early nephron (*Wnt4*), podocyte (*Mafb, Nphs1*), distal tubule (*Slc12a1*), stroma (*Col3a1, Foxd1*), endothelium (*Pecam1*) and macrophages (*Tryobp*).

**Supplementary Figure 3.**
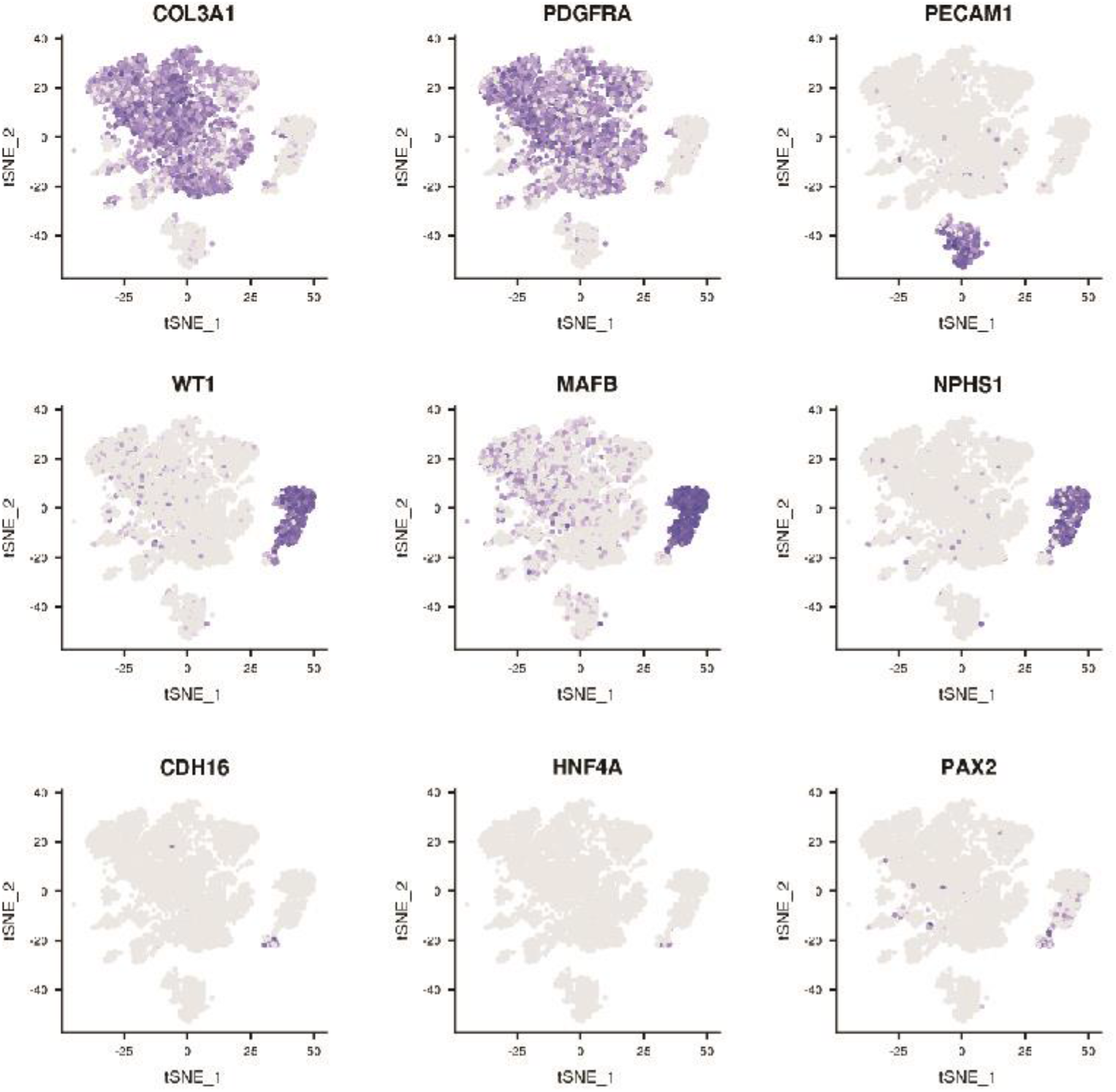
Marker expression across cells in the human iPSC-derived kidney organoid. tSNE plot of all cells from kidney organoids showing the location of cells expressing key marker genes, including markers of stroma (Col3a1, Pdgfra), endothelium (PECAM1), podocyte (WT1, MAFB, NPHS1) and early nephron (CDH16, HNF4A, PAX2).

**Supplementary Figure 4.**
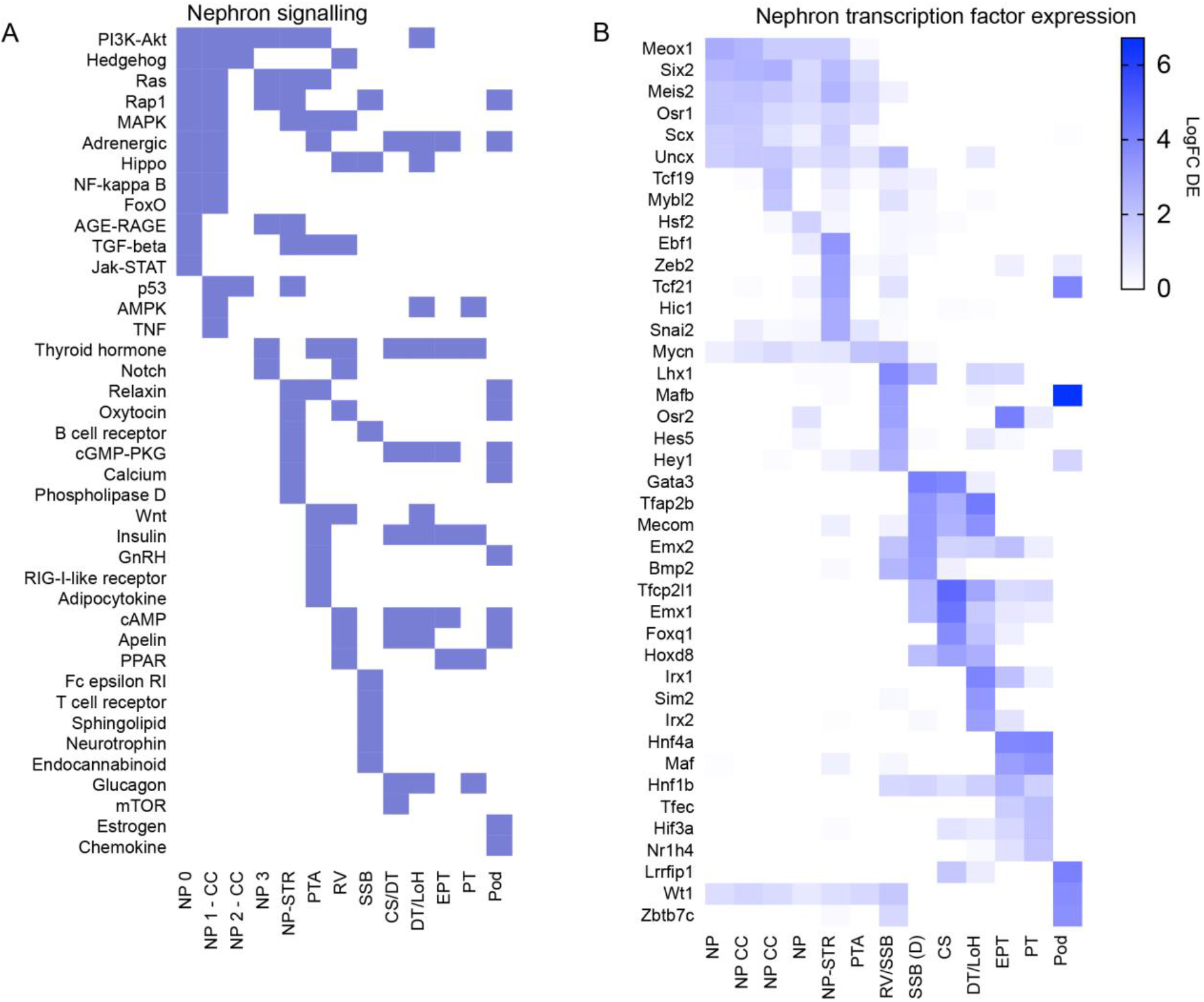
Roadmap of signalling pathways and transcription factros active within the developing mouse kidney. **A.** Transcriptional evidence of signalling pathways active within individual nephron lineage clusters identified by GO and KEGG analysis. Information about which ligands, receptors, and effectors are expressed in each cell type can be accessed in Supplementary table 4. **B**. Expression of key transcription factors within individual nephron lineage clusters.

**Supplementary Figure 5.**
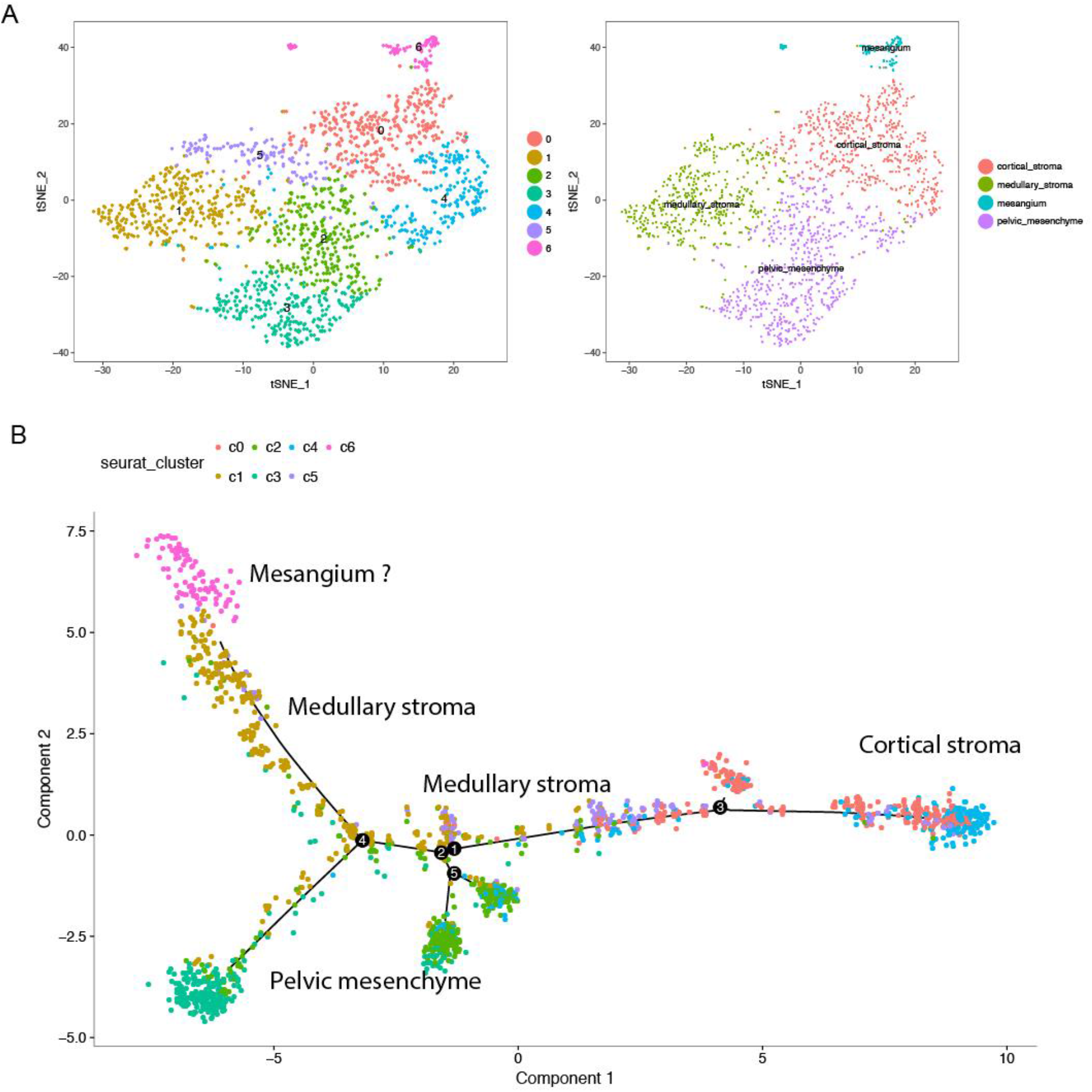
Cluster comparison and Monocle psuedotime clustering of stromal populations from the E18.5 developing mouse kidney. **A**. Comparison of stromal populations identified from clustering the stromal lineage alone (left), to stromal clusters identified within the whole developing kidney (right). **B**. Monocle analysis of all cells within the stromal lineage identifies a trajectory from cortical stroma through intermediate populations to split between pelvic mesenchyme and medullary stroma. The medullary stroma trajectory terminates with a population that may represent mesangial cells.

